# A vibrissa pathway that activates the limbic system

**DOI:** 10.1101/2021.08.02.454720

**Authors:** Michaël Elbaz, Amalia Callado-Pérez, Maxime Demers, Shengli Zhao, Conrad Foo, David Kleinfeld, Martin Deschênes

## Abstract

Vibrissa sensory inputs play a central role in driving rodent behavior. These inputs transit through the sensory trigeminal nuclei, which give rise to the ascending lemniscal and paralemniscal pathways. While lemniscal projections are somatotopically mapped from brain stem to cortex, those of the paralemniscal pathway are more widely distributed. Yet the extent and topography of paralemniscal projections are unknown, along with the potential role of these projections in controlling behavior. Here we used viral tracers to map paralemniscal projections. We find that this pathway broadcasts vibrissa-based sensory signals to brain stem regions that are involved in the regulation of autonomic functions and to forebrain regions that are involved in the expression of emotional reactions. We further provide evidence that GABAergic cells of the Kölliker-Fuse nucleus gate trigeminal sensory input in the paralemniscal pathway via a mechanism of presynaptic or extrasynaptic inhibition.

## INTRODUCTION

Most sensory systems comprise parallel pathways of sensory information that encode different features of a stimulus and also take part in the control of sensor motion (Merigan and Maunsell, 1993; Lomber and Malhotra, 2008; Nassi and Callaway, 2009; Niu et al., 2013; Igarashi et al., 2012). The vibrissa system of rodents is no exception (reviewed in Kleinfeld and Deschênes, 2011). Ascending signaling in the vibrissa system comprises two main trigeminothalamic pathways (reviewed in Deschênes and Urbain, 2016): (1) a lemniscal pathway that arises from the trigeminal nucleus principalis (PrV), transits through the ventral posterior medial nucleus (VPM) of the thalamus, and projects to the primary somatosensory cortex; (2) a paralemniscal pathway that arises from the rostral part of trigeminal nucleus interpolaris (SpVIr), transits through the posterior group (Po) of the thalamus, and projects to the somatosensory cortical areas and to the vibrissa motor cortex.

In contrast with PrV cells, which innervate principally VPM thalamus, SpVIr cells that project to Po thalamus also innervate a number of additional regions by means of branching axons (Pierret et al., 2000; Veinante et al. 2000, Bellavance et al., 2017). These include the superior colliculus, the anterior pretectal nucleus, the ventral division of zona incerta, and the dorsal lateral sector of the facial nucleus. Tract tracing studies by means of classic tracers also reported interpolaris projections to the brain stem and spinal cord (Jacquin et al., 1986; Phelan and Falls, 1991). Whether the latter projections also arise from Po-projecting cells remains an open question.

While the lemniscal pathway conveys tactile information, as well as information about the relative phase of the vibrissae in the whisk cycle (Yu et al., 2006; Curtis and Kleinfeld, 2009; Khatri et al. 2010; Moore et al., 2015; Isett and Feldman, 2020), the role of the paralemniscal pathway remains puzzling. It was proposed that this pathway conveys information about whisking kinematics (Yu et al., 2006), but later studies found that encoding of whisking along the paralemniscal pathway is relatively poor (Moore et al., 2015; Urbain et al., 2015). It was also proposed that the paralemniscal pathway is specifically activated upon noxious stimulation (Masri et al., 2009; Frangeul et al., 2014), but it has never been shown that interpolaris cells that respond to vibrissa deflection are also activated by noxious stimuli. Thus the general function of the paralemnical pathway remains unresolved.

Here we used virus-based tract tracing methods and electrophysiology to document the full extent of the collateral projections of interpolaris cells that give rise to the paralemniscal pathway. We searched for new pathways of ascending information to forebrain regions as well as for feedback projections to brain stem nuclei as a means to uncover the role of the paralemniscal pathway in rat’s behavior.

## RESULTS

### Collateral Projections of Interpolaris Cells that Project to Po Thalamus

To map the collateral projections of interpolaris cells that innervate Po thalamus, we injected the retrograde virus G-pseudotyped Lenti-Cre in Po thalamus, and AAV2/8-hSyn-DIO-GFP in the vibrissa-responsive sector of the SpVIr subnucleus (5 juvenile rats; **Figure 1A**). This approach revealed an unexpected diversity of axonal projections across all animals (**Figure 1B-G**). In accordance with prior studies, terminal fields are found in Po thalamus, the ventral division of zona incerta, the anterior pretectal nucleus, the superior colliculus, the perirubral region, the dorsal lateral part of the facial nucleus, and in the other subdivisions of the sensory trigeminal complex (Jacquin et al., 1986, 1989; Veinante et al., 2000; Bellavance et al., 2017). As new results, profuse projections are also found in the ipsilateral dorsal medullary reticular nucleus (MdD) and further caudally in the dorsal horn of the cervical cord. An ascending projection terminates ipsilaterally in the intertrigeminal region (ITr) and in the Kölliker-Fuse/parabrachial complex (KF/PBc). Whether individual interpolaris cells project to many or all of the above-mentioned target regions remains an open issue. What is clear, however, is that interpolaris cells that project to Po broadcast sensory messages to multiple midbrain and hindbrain regions.

**Figure 1 with 1 supplement.**
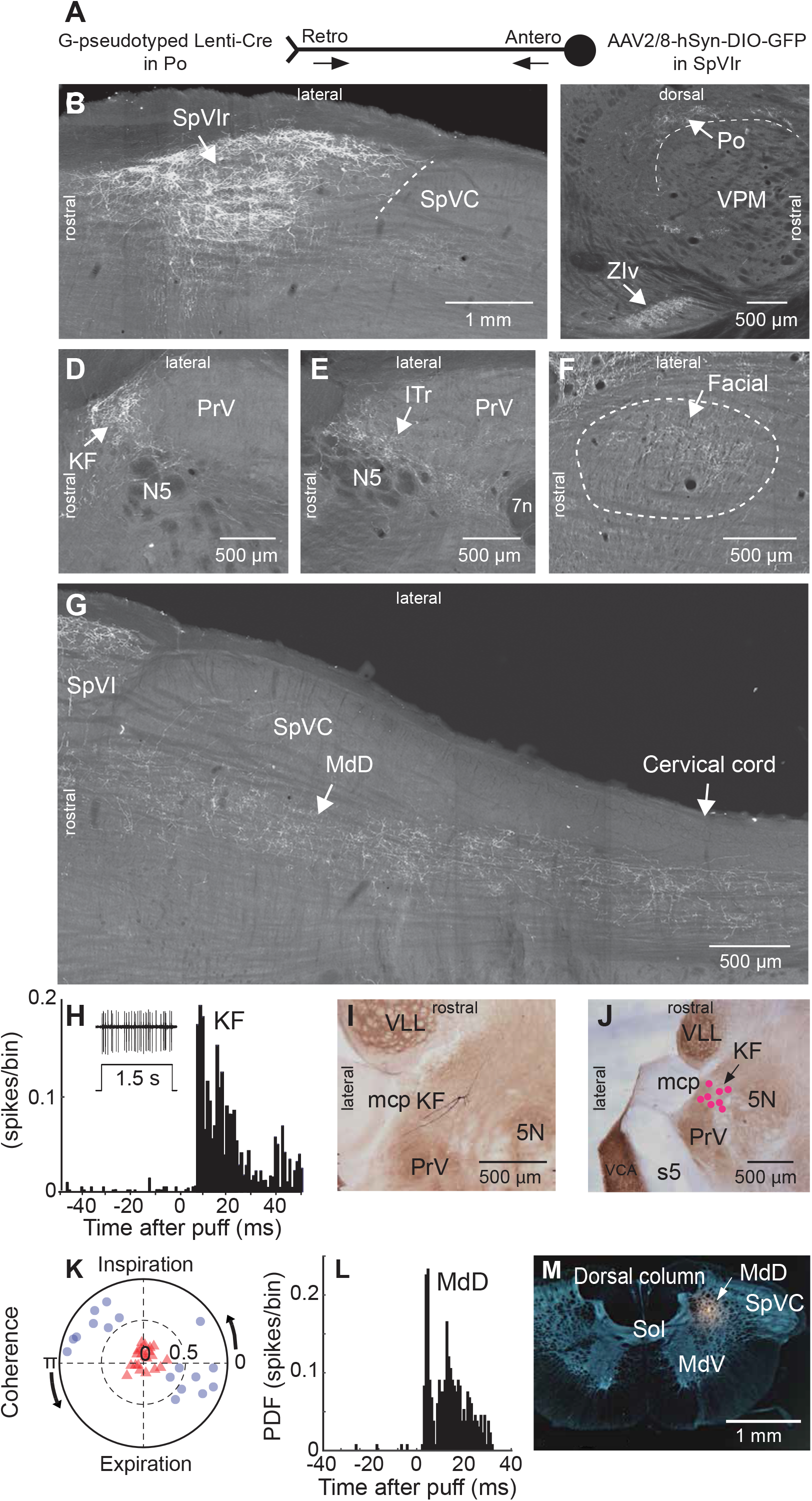
Anatomical and Electrophysiological Evidence that Vibrissa-Responsive Interpolaris Cells Have Widespread Axonal Projections. **(A)** Viral method used for labeling paralemniscal projections. **(B)** Labeling of interpolaris cells after injection of G-pseudotyped Lenti-Cre virus in Po thalamus, and a Cre-dependent AAV that expresses GFP in the vibrissa-responsive sector of SpVIr. **(C)** Anterograde labeling of terminal fields in Po thalamus and zona incerta. **(D)** Anterograde labeling in KF/PB. **(E)** Anterograde labeling in the ITr. **(F)** Anterograde labeling in the dorsal sector of the facial nucleus. **(G)** Anterograde labeling in the MdD and cervical cord. **(H)** Population peristimulus time histogram of spike discharges evoked in KF (22 cells) by air puff deflection of the vibrissae in the anesthetized rat. A representative response is shown in the insert. **(I)** Example of a vibrissa responsive KF cell labeled by juxtacellular delivery of Neurobiotin. **(J)** Location of 8 juxtacellularly labeled KF cells. Horizontal brain stem sections in panels I and J were counterstained for cytochrome oxidase. **(K)** Spectral coherence of spontaneous discharges of KF cells with the respiratory cycle at the respiratory frequency; 1 to 3 Hz. Note that, in contrast with the respiratory units (blue dots), spontaneous discharges of vibrissa-responsive cells (red triangles) display low coherence with respiration. **(L)** Population peristimulus time histogram of spike discharges evoked in MdD (33 cells) by air puff deflection of the vibrissae. **(M)** Recording site in the MdD labeled by an iontophoretic injection of Chicago Sky Blue. This coronal section was counterstained for cytochrome oxidase and a negative image was generated. See **Figure S1** for additional anatomical data. **Abbreviations for all anatomy:** 5n, root of the trigeminal motor nucleus; 5N, trigeminal motor nucleus; 5t, trigeminal tract; 7n, facial nerve tract; 7N, facial nucleus; Amb, ambiguus nucleus; APT, anterior pretectal nucleus; BSTL, bed nucleus of the stria terminalis; CeA, central amygdala; Cerv Cord, cervical cord; CM/PC, central medial/paracentral thalamic nuclei; CPu, caudate putamen; DR, dorsal raphe; EW, Edinger-Westphal; Hab, habenula; IML, intermedio-lateral column of the spinal cord; ITr, intertrigeminal region; KF, Kölliker-Fuse nucleus; KF/PB, Kölliker-Fuse/parabrachial complex; mcp, middle cerebellar peduncle; MdD, dorsal part of the medullary reticular formation; MdV, ventral part of the medullary reticular formation; mt, mammillothalamic tract; NA, nucleus ammbiguus; NTS, nucleus of the solitary tract; opt, optic tract; PAG, periaqueductal gray; PB, parabrachial nuclei; PC, paracentral thalamic nucleus; PCRt, parvicellular reticular formation; PLH, posterior lateral hypothalamus; Po, posterior nuclear group of the thalamus; PrV, principal trigeminal nucleus; RN, red nucleus; Rt, reticular thalamic nucleus; s5, sensory root of the trigeminal nerve; SC, superior colliculus; scp, superior cerebellar peduncle; SpVC, caudalis division of the spinal trigeminal complex; Sol, nucleus of the solitary tract; SpVIc, caudal sector of the interpolaris trigeminal nucleus; SpVIr, rostral division of the interpolaris nucleus; TG, trigeminal ganglion; VLL, ventral nucleus of the lateral lemniscus; VPL, ventral posterolateral thalamic nucleus; VPM, ventral posterior medial nucleus; VPCc, parvicellular sector of the ventral posteromedial thalamic nucleus; VPPc, parvocellular part of the ventral posterior thalamic nucleus; VRG, ventral respiratory group; vsc, ventral spinocerebellar tract; ZIv, ventral division of zona incerta.

Injection of the anterograde virus AAV1-hSyn-eGFP-WPRE in the vibrissa-responsive sector of the SpVIr (3 rats; **Figure 1 - supplemental figure 1**) yielded similar results to those above (**Figure 1B G**); yet in these cases anterograde labeling is also found in the cerebellum. This is consistent with prior studies, which reported that different populations of SpVI cells project to the thalamus and cerebellum (Steindler, 1985; Jacquin et al., 1986). Of note, in sagittal sections of the brain stem, the terminal field of interpolaris axons in KF/PBc covers an extensive crescent-shaped territory at the rostral border of PrV and the trigeminal motor nucleus (**Figure 1 - supplemental figure 1**).

### KF/PBc and MdD Cells Respond to Vibrissa Deflection

Interpolaris cells that give rise to the paralemniscal pathway respond to vibrissa deflection (Veinante et al., 2000). Thus, one expects KF/PBc and MdD neurons to also respond to vibrissa deflection. We indeed found vibrissa-responsive neurons in the KF/PBc (latency: 8.0 ± 1.2 ms; mean ± SD; 22 cells across 14 animals; **Figure 1H K**) and in the MdD (latency: 6.0 ± 1.4 ms; 33 cells across 3 rats; **Figure 1L,M**) of ketamine/xylazine anesthetized animals. The location of recorded units was assessed by single cell juxtacellular labeling in the KF/PBc (8 cells in 8 of the 14 rats), and by iontophoretic deposit of Chicago Sky Blue in the MdD (3 rats). Like interpolaris cells, the receptive field of KF/PBc and MdD cells includes multiple vibrissae. In KF/PBc, vibrissa-responsive cells are intermingled with cells that discharge in phase with the respiratory rhythm. Yet the spontaneous activity of vibrissa-responsive cells displays no statistically significant spectral coherence with respiration (mean coherence <C> in the spectral band of breathing is |<C>| = 0.16 for vibrissa-responsive cells versus |<C>| = 0.72 for respiratory units (**Figure 1K**).

### Axonal Projections of KF/PBc Cells that Receive Interpolaris Inputs

Among the targets of the paralemniscal pathway, we focused on the KF/PBc because of the well-documented hodology of its axonal projections (see review by Saper, 2015). As a means to visualize the axonal projections of KF/PBc cells that receive interpolaris input, we injected AAV2/1 that expresses Cre in the vibrissa-responsive sector of the SpVI to achieve anterograde transsynaptic expression of Cre (Zingg et al., 2017), and AAV1-EF1a-DIO-hChR2-eYFP was injected in the KF/PBc (3 rats). Transsynaptically labeled cells amount to 55, 62, and 68 in each of the rats. We observed transsynaptic labeling anterograde labeling in the lateral part of the facial nucleus, throughout the ventral lateral medulla, the MdD, the nucleus of the solitary tract, and the cervical cord in all animals (**Figure 2A-D**). Ascending projections were found in the dorsal raphe, the Edinger-Wesphal nucleus, the periaqueductal gray, the posterior lateral hypothalamus, the paracentral and central medial thalamic nuclei, the parvicellular part of the ventral posterior thalamic nucleus, the ventral division of zona incerta, the central amygdala (CeA), and the lateral part of the bed nucleus of stria terminalis in all animals (**Figure 2E-G and Figure 2 - supplemental figure 1**). In summary, KF/PBc cells that receive interpolaris input in turn project to most of the brain stem and forebrain regions previously identified by means of anterograde tracer injections in the KF/PBc (Saper, 2015) (**Figure 2H)**. Yet, it is worth noting that no anterograde labeling is observed in the trigeminal sensory nuclei. This negative result is of critical importance. It indicates that KF/PB cells that receive interpolaris input do not feed back to the relay cells that give rise to the paralemniscal pathway (see below).

**Figure 2 with 1 supplement.**
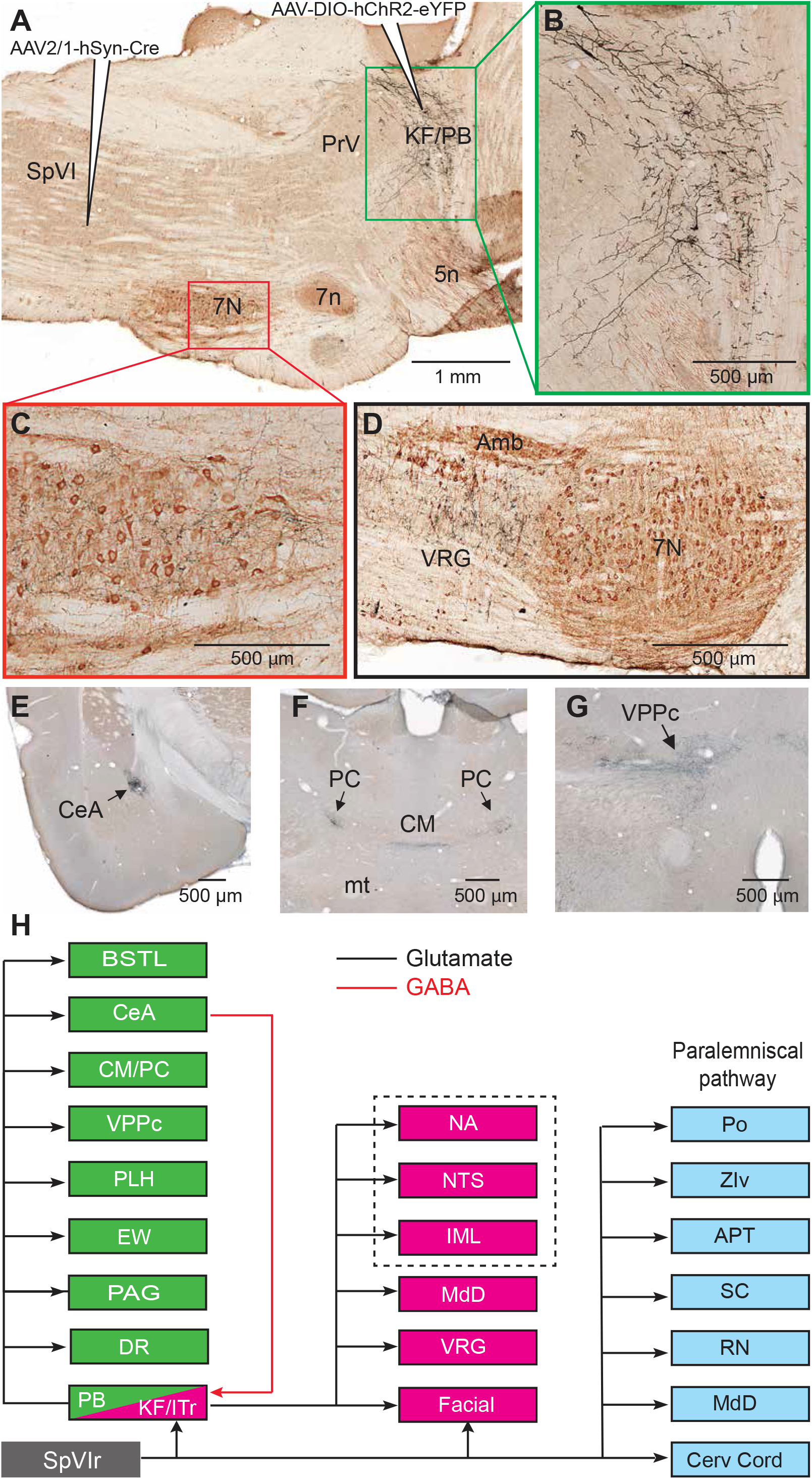
Axonal Projections of KF/PB Cells that Receive Interpolaris Input. See **Figure 1** for abbreviations. **(A-C)** AAV2/1-hSyn-Cre-WPRE was injected in the vibrissa-responsive sector of the SpVIr, and AAV2/1-EF1a-DIO-hChR2-eYFP was injected in KF/PB. This parasagittal section shows labeling in KF/PB. The section was immunostained for choline acetyltransferase (A). The green and red framed areas are enlarged in (B) and (C). **(D)** Terminal labeling in the ventral respiratory group underneath the ambiguus nucleus. **(E)** Anterograde labeling in the central amygdala. **(F)** Anterograde labeling in the paracentral and central medial thalamic nuclei. **(G)** Anterograde labeling in the parvocellular division of the ventral posterior medial nucleus of the thalamus (see **Figure S2** for additional projection sites). **(H)** Summary of the first-order axonal projections of interpolaris cells, and the second-order projections that derive from the KF/PB complex. Projection sites in the framed area are from prior studies (see review by Saper and Stornetta, 2014).

As PB cells that project to the CeA receive input from vibrissa-responsive SpVI neurons, one expects to find vibrissa-evoked responses in the amygdala. We addressed this issue by recording the local field potential evoked by air jet deflection of the vibrissae (3 rats; **Figure 3A C**). These experiments were carried out under urethane instead of ketamine/xylazine anesthesia as α2 agonists, e.g., xylazine, strongly depress synaptic transmission in the parabrachial-amygdaloid pathway (Delaney et al., 2007). Vibrissa deflection evoked a clear negative field potential in the CeA (peak latency of 12 ms as an average over all 3 rats and 50 trials per rat; **Figure 3A,B**). This response was abolished in all animals following electrolytic lesion of the ipsilateral PB (**Figure 3A,C**). Together, our anatomical (**Figure 2**) and electrophysiological (**Figure 3A-C**) data delineate a pathway of vibrissa information that reaches the CeA and several limbic regions of the forebrain via a relay in the KF/PBc.

**Figure 3 with 1 supplement.**
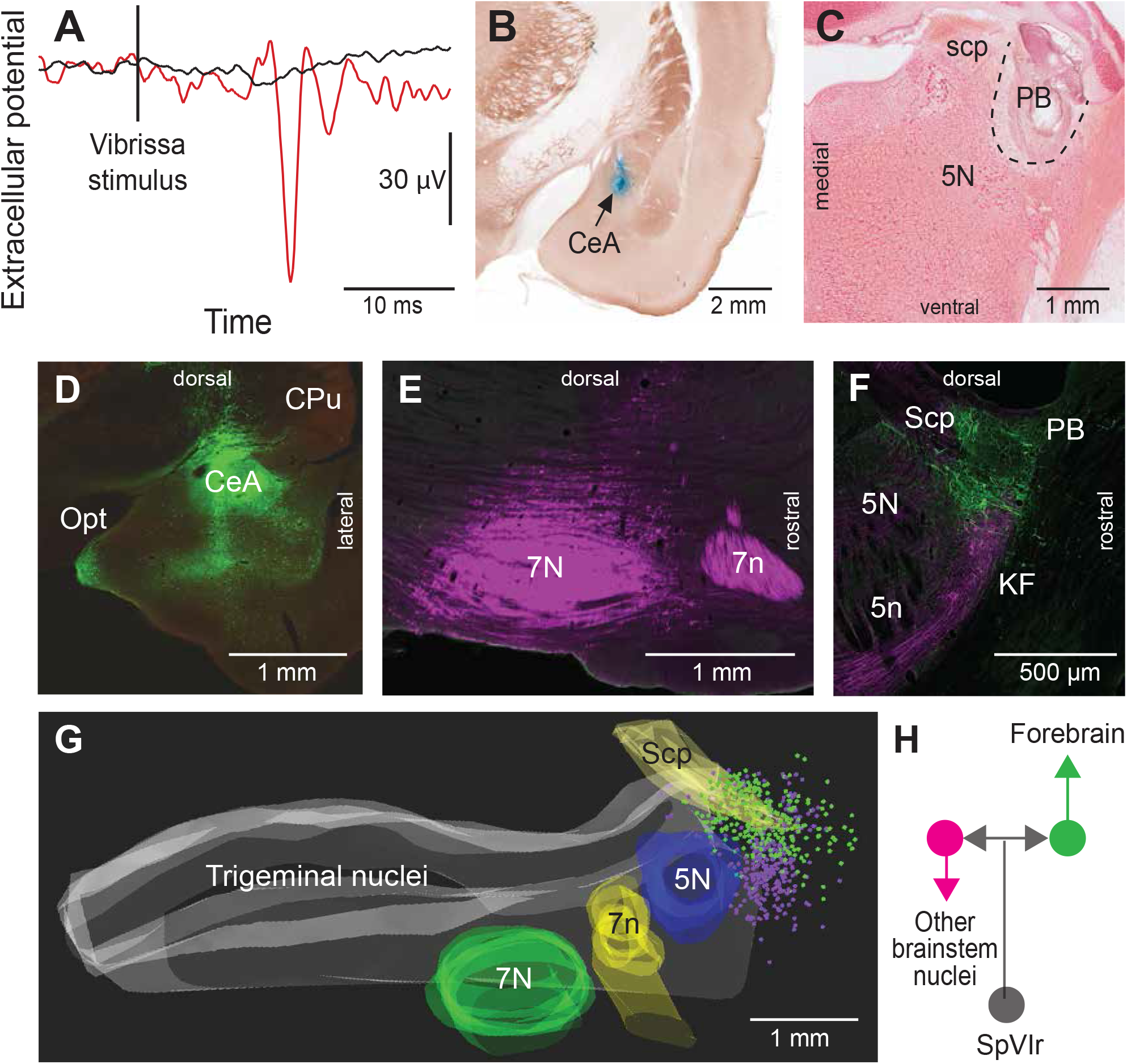
Vibrissa-Evoked Responses in Central Amygdala and Anatomical Evidence that Separate Cellular Populations in KF/PB Project to the CeA as Compared to the Facial Nucleus. See **Figure 1** for abbreviations. **(A)** Average response (50 trials) evoked in the CeA by air puff stimulation of the vibrissae before (red trace) and after (black trace) an electrolytic lesion of the PB complex. **(B)** Recording site in the amygdala labeled by an iontophoretic injection of Chicago Sky Blue (cytochrome oxidase counterstaining). **(C)** Electrolytic lesion of the PB complex. **(D)** Injection site of retroAAV-CAG-eGFP in the CeA. **(E)** Injection site of retroAAV-CAG-mCherry in the facial nucleus. **(F)** Retrogradely labeled cells in KF/PB. **(G)** 3D reconstruction showing the distribution of amygdala-projecting cells (green dots) in the medial and lateral PB, and facial-projecting neurons in the KF (magenta dots). Note that a few cells in lateral PB project to the facial nucleus; yet, none of the KF/PB cells is doubly labeled. **(H)** Wiring diagram of the projections of KF and PB cells that receive vibrissa input from the interpolaris nucleus (see also **Figure S3** for additional evidence).

### Differential Projections of KF and PB Cells

Prior studies in mice have shown that separate pools of KF/PB neurons project to the forebrain and to lower brain stem (Greeling et al., 2017; Barik et al., 2018). We thus examined whether descending and ascending projections of KF/PBc in rats arise from different or overlapping neurons. We made large injections (100 nl) of the retrograde labels retroAAV-CAG-eGFP and retroAAV-CAG-mCherry in the CeA and the facial nucleus, respectively (2 rats; **Figure 3D-G**). Although a number of retrogradely labeled cells were found in the KF/PBc, none was doubly labeled. In detail, amygdala-projecting cells (326 in rat 1 and 312 in rat 2) were mainly concentrated in lateral and medial PB, while the majority of facial projecting cells (299 in rat 1, and 140 in rat 2) were located in KF. In complementary experiments, we injected retroAAV expressing Cre in the ventral respiratory group and AAV2/8-hSyn-DIO-eGFP in KF/PBc (2 rats). Anterograde labeling was observed in the facial nucleus, the ventral respiratory group and the MdD (**Figure 3 - supplemental figure 1**), but not in midbrain and forebrain regions. Together these results confirm that neighboring but separate populations of KF/PB neurons project to the forebrain and to the lower brain stem (**Figure 3H**).

### GABAergic KF cells gate sensory transmission in the SpVI

In rodents, KF projections to the brain stem arises from two neuronal populations: glutamatergic cells that project to the autonomic, respiratory, and motor regions of the medulla, and GABAergic neurons, which project principally to the SpVI (Geerling et al., 2017). To determine whether GABAergic KF cells inhibit interpolaris cells, we made large injections (200 nl) of the transsynaptic label anteroAAV-expressing Cre (Zingg et al., 2017) in KF/PB and AAV2-Ef1a-DIO-eYFP in the SpVI (3 rats; **Figure 4A-D**). Unexpectedly, few interpolaris neurons were labeled (44, 48, and 53 cells respectively in each of the rats). Anterograde labeling was observed in the KF/PB complex and in the parvicellular part of the ventral posterior thalamic nucleus, a region known to process input from the tongue and oral cavity (Ogawa and Nomura, 1988; Verhagen et al., 2003), in all animals. No labeling was observed in the cerebellum, or in brain stem and midbrain regions that receive input from the paralemniscal pathway (**Figure 2H**). Yet, cells in the parvicellular reticular formation (PCRt) at the medial border of the trigeminal nuclei were extensively labeled, which indicates that the paucity of cell labeling in SpVI is hardly ascribable to a technical pitfall. Together these results indicate that KF projections do not innervate the SpVI interneurons or the projection cells that give rise to the paralemniscal pathway.

**Figure 4.**
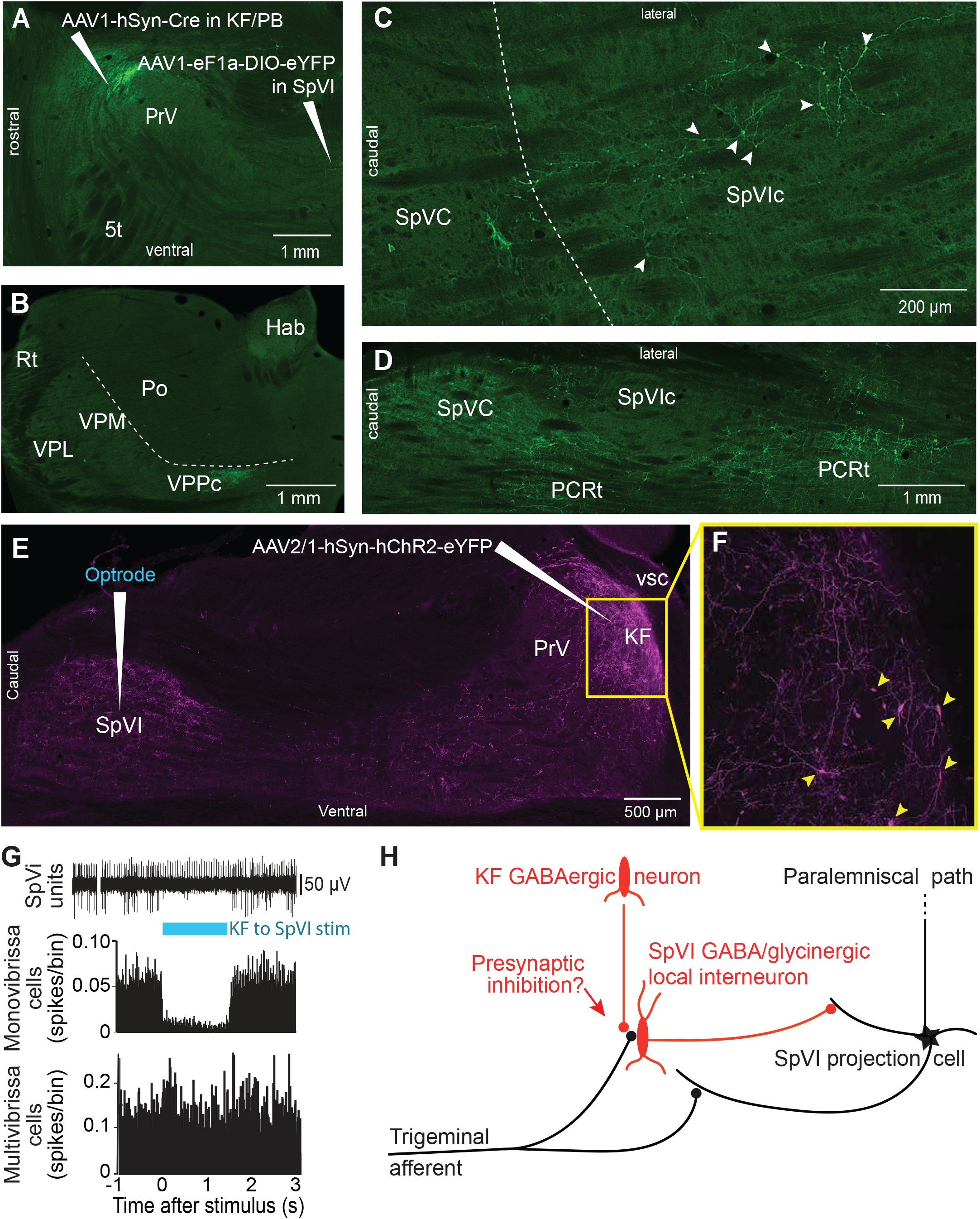
Postsynaptic targets of KFn projections to the SPVI. See **Figure 1** for abbreviations. (**A**) The viral method used for transsynaptic labeling of brain stem cells that receive input from the KFn. (**B-D**) Transsynaptic labeling in SpVI is restricted to a small population of cells (C) that project to VPCC (B). Yet, numerous cells are transsynaptically labeled in the parvicellular reticular formation (D). (**E,F**) Viral labeling method for optogenetic stimulation of KFn axons. Injection of AAV2/1-hSyn-hChR2-eYFP in KF results in anterograde labeling in SpVI. The framed region in E is shown in F; arrowheads point to labeled cell bodies. Note the dense network of axon collaterals within the KF/PB complex. **(G)** Representative responses of a monovibrissa-responsive cell (biphasic unit) and a multivibrissa-responsive cell (positive unit) upon optogenetic stimulation of KF axons labeled as in panel E. Population peristimulus time histogram shows the responses of 26 monovibrissa-responsive units upon optogenetic stimulation of KF axons along with 10 multivibrissa-responsive units to optogenetic stimulation of KF axons. Only mono-vibrissa and not multi-vibrissa cells are responsive to KFn input. **(H)** Summary of the effect of KFn projections on the flow of sensory input through SpVI.

To further assess these anatomical results, we injected AAV-hSyn-ChR2-eYFP in KF and used an optrode to concurrently stimulate KF axons and record the activity of SpVI neurons (3 rats; **Figure 4E-G**). Interpolaris cells either discharged spontaneously or were driven by jiggling a single vibrissa with a piezoelectric stimulator. Optogenetic stimulation suppressed sensory-evoked discharges in monovibrissa-responsive cells (26 cells in 3 rats, t-test yields p < 0.001; lower panel in **Figure 4G**) but did not affect the firing of multivibrissa-responsive units (32 cells in the same 3 rats, p = 0.84; **Figure 4G**). Together with the lack of transsynaptic labeling of local circuit cells from the KF, these results raise the possibility that KF-induced suppression of sensory responses in local circuit interpolaris cells is mediated by presynaptic or extrasynaptic inhibition, as summarized in **Figure 4H**.

## DISCUSSION

Our study reveals a hitherto unsuspected diversity of axonal projections of interpolaris cells that give rise to the paralemniscal pathway. We identified two new collateral projections that target the MdD and the KF/PBc. Furthermore, we show that the KF/PBc contains two populations of vibrissa-responsive neurons; one gives rise to an ascending pathway, which projects to limbic regions of the forebrain, and the other one projects to the autonomic, respiratory, and motor regions of the medulla. Lastly, KF contains a pool of GABAergic neurons, which do not receive interpolaris input, but project locally and to the trigeminal sensory nuclei. The diagram in **Figure 2H** summarizes the first-order axonal projections of interpolaris cells that give rise to the paralemniscal pathway, and the second-order projections that derive from the KF/PBc.

### Descending Projection to the MdD

By far the most abundant collateral projection of vibrissa-responsive interpolaris cells is to the MdD and the dorsal horn of the cervical cord. The finding of vibrissa-responsive cells in the MdD is unexpected as most studies reported that MdD neurons best respond to noxious stimuli (reviewed in Martins and Tavares, 2017). Although there is no reason to believe that vibrissa deflection per se is painful, an unexpected air puff directed toward the head of a rat elicits fear-related behavior such as a startle response (Engelmann et al., 1996), 22 kHz ultrasonic vocalization (Knapp and Pohorecky, 1995) and avoidance behavior (Cimadevilla et al., 2001). As the MdD sends profuse projections to the cervical cord and the facial nucleus (Bernard et al., 1990; Takatoh et al., 2013; Takatoh et al., 2021), these results suggest that vibrissa messages to the MdD may elicit avoidance or flight behaviors. This conclusion is in line with that of the recent study, which reported that the MdD is part of a brain stem-spinal circuit to control escape responses to noxious stimuli (Barik et al., 2018).

### The Trigemino-Parabrachial Amygdaloid Pathway

A number of studies have reported trigeminal projections to the KF and PB subnuclei (Cechetto et al., 1985; Slugg and Light 1994; Feil and Herbert, 1995; Dallel et al., 2004; Rodriguez et al., 2017; see also review by Chamberlin, 2004). Most of these studies focused on ascending projections from subnucleus caudalis, which convey pruriceptive and nociceptive inputs to the KF/PBc (Jansen and Giesler, 2015). Thus, it came as a surprise to find vibrissa-responsive neurons in KF/PBc, and further find that vibrissa signals are conveyed to the limbic regions of the forebrain via the medial and lateral PB. One study previously reported short-latency activation of amygdala neurons, at about 11 ms, upon electrical stimulation of the mystacial pad (Bernard et al., 1992). Such a short latency is consistent with the activation of a trigemino-parabrachial-amygdaloid pathway by Aβ primary vibrissa afferents.

### The Paralemniscal Pathway and Threat

Although vibrissa-responsive interpolaris cells do not respond to noxious stimuli, many of the regions they innervate contain cells that process nociceptive or aversive inputs. This is the case for the MdD (reviewed in Martins and Tavares, 2017), the KF/PBc (Jansen and Giesler, 2015), the superior colliculus (Dean et al., 1989; Redgrave et al., 1996), the zona incerta (Masri et al., 2009), and Po thalamus (Masri et al., 2009; Frangeul et al., 2014; Sobolewski et al., 2015). These anatomical data suggest that the paralemniscal pathway conveys signals from threatening or alarming sensory inputs, perhaps through association with aversive contexts.

In light of the evidence that interpolaris cells innervate brain regions that process nociceptive or aversive stimuli, we postulate that the KF is primarily involved in orchestrating defensive reactions to aversive and threatening stimuli. Three additional lines of evidence support this hypothesis. First, glutamatergic KF cells project to the cardio-respiratory medullary centers, the facial, hypoglossal and ambiguous motor nuclei, the nucleus of the solitary tract, and to the preganglionic neurons of the sympathetic nervous systems (reviewed in Saper and Stornetta, 2015). Second, activation of KF cells elicits changes in respiration, heart rate, blood pressure (Chamberlin and Saper, 1992; Lara et al., 1994; Guo et al., 2002; Chamberlin, 2004). Third, individual KF neurons project to many of the above-mentioned targets by means of branching axons, which might form the anatomical substrate for coordinating autonomic and somatic reactions (Song et al, 2012).

### Sensory Gating in the Paralemniscal Pathway

A prior study in the alert head-restrained rat reported that units in trigeminal nucleus principalis fire more spikes when the animal is whisking as opposed to not whisking. Yet spike rates are not significantly different between whisking and not whisking in nucleus SpVI (Moore et al., 2015). This is an unexpected result, as primary vibrissa afferents discharge during whisking (Severson et al., 2017) and SpVI projection cells are typically driven by many more vibrissae that principalis neurons (Furuta et al., 2006). The observation that whisking tends to reorganize spike times of neurons in interpolaris, rather than increase overall spike rates, supports the notion that the relay of vibrissa messages in the paralemniscal pathway is normally gated. The inhibition of the paralemniscal pathway by Kölliker-Fuse GABAergic neurons, together with the lack of a direct connection from Kölliker-Fuse to interpolaris projections cells, suggests that gating occurs by a mechanism of presynaptic or extrasynaptic inhibition (**Figure 4H**).

## Conclusion

Our results clearly indicate that the paralemniscal pathway broadcasts vibrissa-based sensory signals to a number of brain stem and forebrain regions that are involved in the expression of emotional reactions. They provide a different viewpoint on the role of the Kölliker-Fuse nucleus, which is currently considered to control the mechanics of respiratory and cardiac rhythms, but no more. We propose that the Kölliker-Fuse nucleus is involved in the larger role of orchestrating the expression of emotional reactions, e.g., facial expressions, autonomic and somatic reactions, of which respiration is one of many players.

## ACKNOWLEDGMENTS

We thank Fan Wang for critical discussions and for supplying the retrograde-Lenti-Cre virus. This work was supported by grants from the Canadian Institutes of Health Research (grant MT-5877) and the National Institutes of Health (U19 NS107466 and R35 NS097265).

## AUTHOR CONTRIBUTIONS

D.K. and M. Deschênes supervised the project. M.E. and A.C-P. performed the experiments. S.Z. prepared reagents. M.E., C.F. and M. Demers analyzed data. D.K. and M. Deschênes wrote the manuscript.

## FIGURE LEGENDS

**Figure 1 supplement 1.**
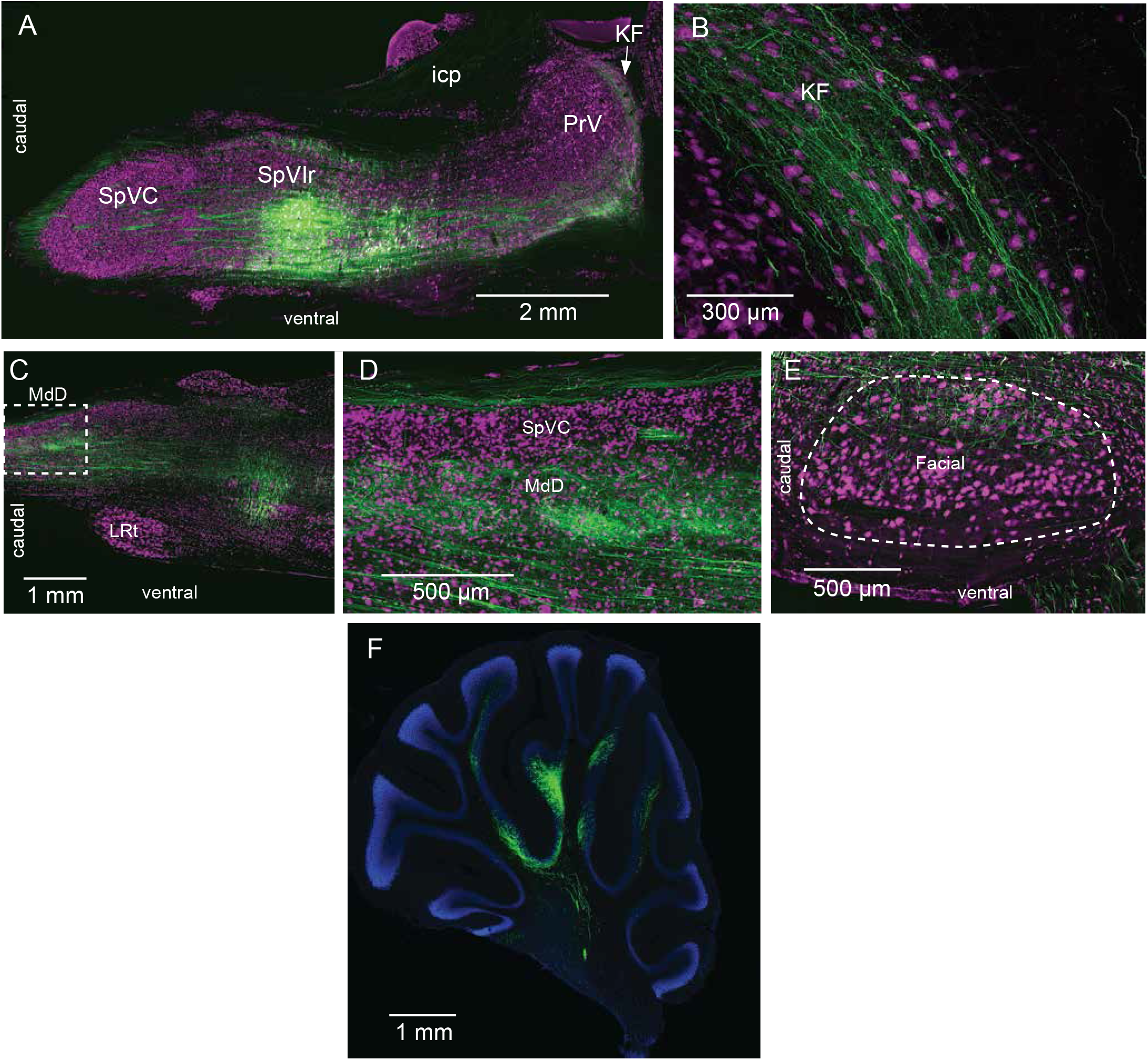
Interpolaris cells project to KF/PB and MdD (Supplementary information related to Figure 1). **(A)** Injection site of AAV1-hSyn-eGFP-WPRE-hgh in SpVIr. **(B)** Anterograde labeling in KF/PB. **(C)** Anterograde labeling in the MdD. The framed region is enlarged in (D). **(E)** Terminal field in the dorsal lateral sector of the facial nucleus. **(F)** Anterograde labeling in the cerebellum. Sections were immunostained for NeuN (A-E), or counterstained with DAPI (F). **Abbreviations**: icp, inferior cerebellar peduncle; KF, Kölliker-Fuse nucleus; LRT, lateral reticular nucleus; MdD, dorsal part of the medullary reticular formation; PrV, principal trigeminal sensory nucleus; SpIr, rostral sector of the interpolaris trigeminal nucleus; SpVC, caudalis division of the spinal trigeminal complex.

**Figure 2 supplement 1.**
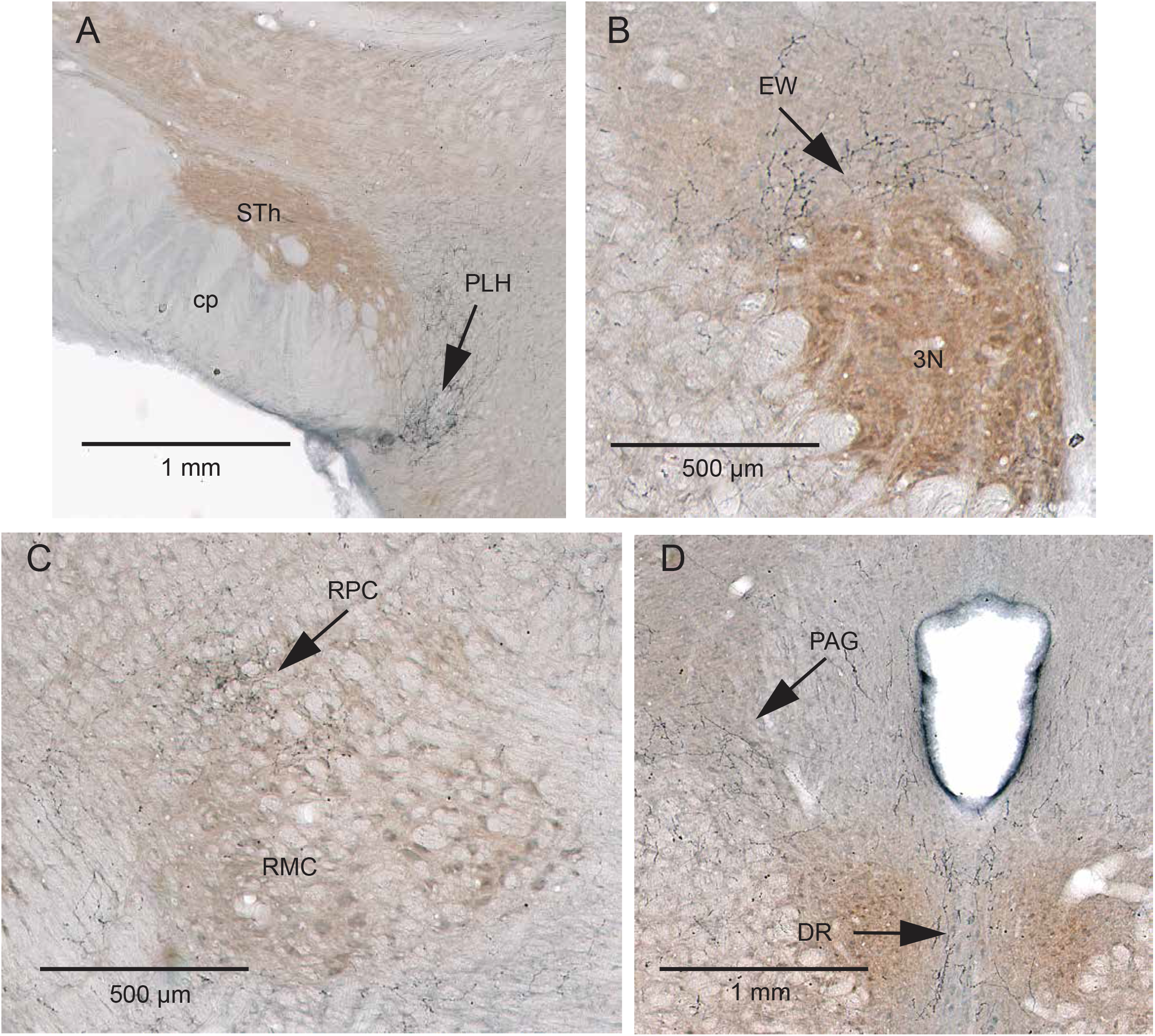
Additional projection sites of KF/PB cells that receive interpolaris input (Supplementary information related to Figure 2). (**A-D**) AAV2/1-hSyn-Cre-WPRE was injected in the vibrissa-responsive sector of the SpVIr, and AAV2/1-EF1a-DIO-hChR2-eYFP was injected in KF/PB. Anterograde labeling is present in the posterior lateral hypothalamus (A), the Edinger-Westphal nucleus (B), the parvocellular division of the red nucleus (C), and in the dorsal raphe and ventral lateral periaqueductal gray (D). Sections were counterstained for cytochrome oxidase. **Abbreviations:** 3N, oculomotor nucleus; cp, cerebral peduncle; DR, dorsal raphe; EW, Edinger-Westphal nucleus; PAG, periaqueductal gray; PLH, posterior lateral hypothalamus; RMC, red nucleus magnocellular part; RPC, red nucleus parvocellular part; STh, subthalamic nucleus.

**Figure 3 supplement 1.**
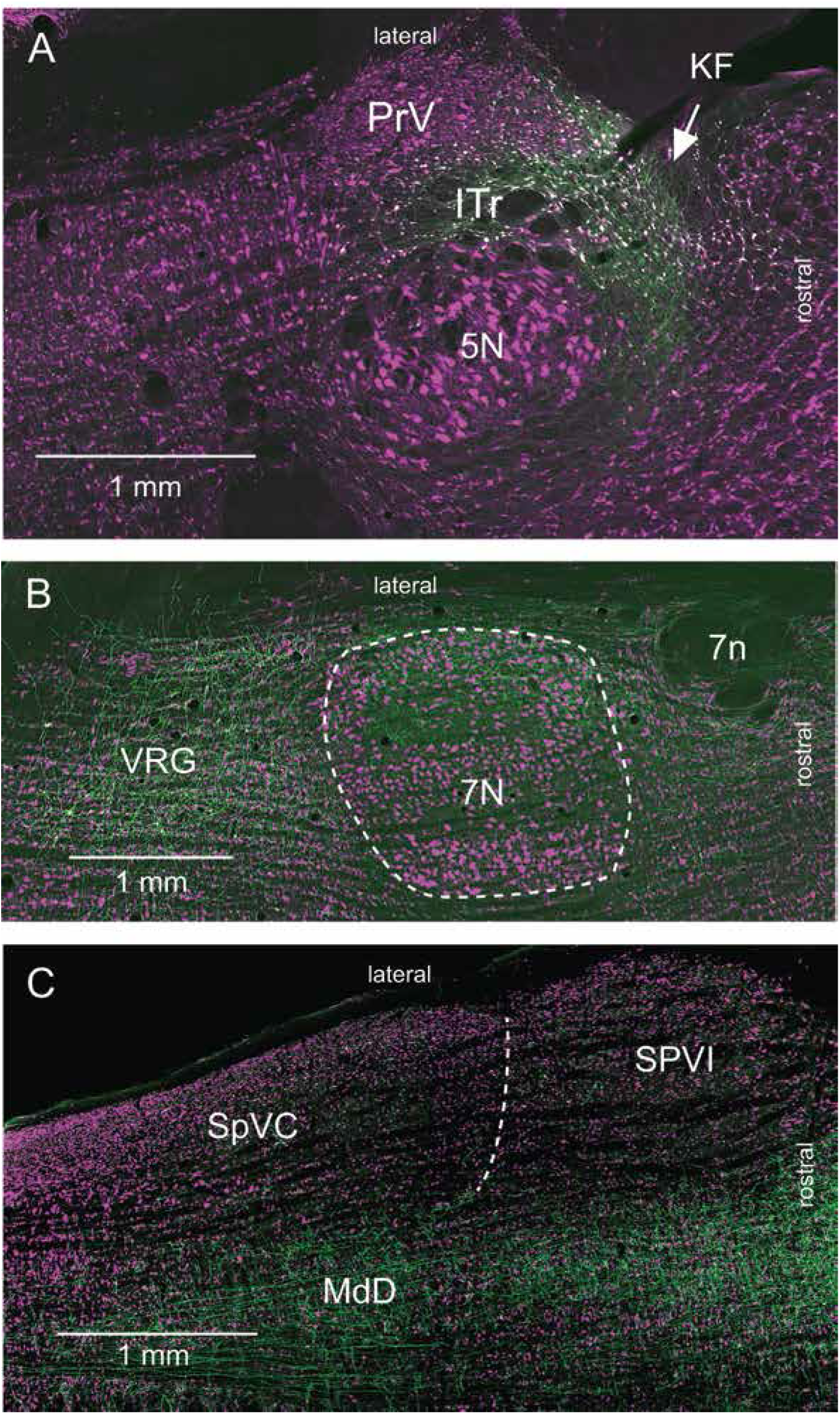
Labeling of KF cells that project to lower brain stem (Supplementary information related to Figure 3). **(A)** Labeling of KF cells after injection of retroAAV-hSyn-Cre in the ventral respiratory group, and AAV2/8-hSyn-DIO-GFP in the KF/PBc. Note the absence of anterograde labeling in the sensory trigeminal nuclei. **(B)** Anterograde labeling in the lateral part of the facial nucleus and in the ventral respiratory group. No anterograde labeling is observed in midbrain or forebrain regions. Horizontal sections were immunostained for NeuN. **Abbreviations:** 5N, trigeminal motor nucleus; 7N, facial nucleus; 7n, facial nerve tract; Amb, ambiguous nucleus; ITr, intertrigeminal region; PrV, principal trigeminal sensory nucleus; SpVC, caudalis division of the spinal trigeminal complex. SpVI, interpolaris division of the spinal trigeminal complex; VRG, ventral respiratory group.

## MATERIALS AND METHODS

### Subjects

Experiments were carried out in Long Evans rats of both sexes and included 5 juvenile rats (P30 - P35) and 34 adult rats (P80 - P100; 250-350 g in mass). All experiments were carried out according to the National Institutes of Health Guidelines. All experiments were approved by the Institutional Animal Care and Use Committees at Laval University and at the University of California at San Diego.

### Viruses

G-pseudotyped Lenti-Cre virus was designed and produced as described (Stanek et al., 2016). AAV2/8-hSyn-DIO-GFP and AAV2/8-hSyn-DIO-GFP were obtained from the Neurophotonics Centre, Laval University. AAV2/1-EF1a-DIO-hChR2-EYFP-WPRE-HGHpA, AAV2/1-hSyn-Cre WPRE, AAV1-hSyn-eGFP-WPRE, retroAAV-CAG-mCherry, retroAAV-CAG-eGFP, AAV2/8-hSyn-DIO-eGFP, and AAV2/8-hSyn-DIO-eGFP were obtained from Addgene.

### Surgery and virus injections

All but three rats were anesthetized with ketamine (75 mg/kg) and xylazine (5 mg/kg), and body temperature was maintained at 37 C with a thermostatically controlled heating pad. Three rats were anesthetized with urethane (1.4 g/kg) to record the local field potential evoked in the amygdala by vibrissa deflection. All virus injections were carried out after electrophysiolological identification of the target regions. When AAV injections were made in the interpolaris nucleus, we targeted the lateralmost sector of the nucleus based on the well-known somatotopic representation of the vibrissae in this nucleus. This precaution proved useful to avoid infecting cells in neighboring regions. After 3 to 4 weeks of survival animals were either perfused with saline and paraformaldehyde (4 % (w/v) in PBS), or anesthetized with ketamine (75 mg/kg) and xylazine (5 mg/kg) for optogenetic stimulation.

In three rats the vibrissa-responsive sector of the KF/PB was first located by electrophysiological recording, and then lesioned by passing 0.5 mA through a stainless steel electrode with a tip (10 µm) de-insulated over a length of 500 µm.

### Electrophysiology

Single unit recordings were carried out in KF/PB and MdD with micropipettes (tip diameter, 1 µm) filled with either 0.5 M potassium acetate and 2 % (w/v) Neurobiotin, or 0.5 M potassium acetate and 4 % (w/v) Chicago Sky Blue. To maximize the chance of cell recovery after juxtacellular labeling in the KF/PB, we injected one cell per animal and perfused the rat thereafter. Vibrissae were deflected in the dorsocaudal direction with air-puffs. The time delay between the command voltage and the onset of vibrissa deflection was measured with a piezoelectric film positioned at the same distance from the puffer tip. Respiration was monitored with a cantilevered piezoelectric film (LDT1 028K; Measurement

Specialties) resting on the rat’s abdomen just caudal to the torso. All signals were sampled at 10 kHz and logged on a computer using the Labchart acquisition system (AD Instruments).

### Optogenetics

Three weeks after injection of AAV2/1-hSyn-hChR2-eYFP in KF/PB (n = 3 rats) we used a carbon-tip optrode (Kation Scientific) to simultaneously stimulate labeled axons and record interpolaris cells. As a test for KF-induced inhibition we drove background spiking by jiggling a single vibrissa with a piezoelectric stimulator.

### Histology

Following perfusion brains were postfixed for 1 h, and cryoprotected overnight in 30 % (w/w) sucrose in PBS. Brains were cut at thickness of 50 µm on a freezing microtome. Labeled material was processed for either fluorescence or brightfield microscopy. For fluorescence microscopy, sections were immunoreacted with a chicken anti-GFP antibody (1:1000; Abcam), and a donkey anti-chicken Alexa 488 IgG (Abcam). NeuN immunostaining was carried out with a rabbit anti-NeuN antibody (1:1000; EM Millipore) and an anti-rabbit IgG conjugated to Alexa 594. For brightfield microscopy, sections were first counterstained for cytochrome oxidase, and then immunoreacted with a rabbit anti-GFP antibody (1:1000; Novus Biological), a biotinylated anti-rabbit IgG (1:200: Vector Labs), the avidin/biotin complex (Vectastain ABC kit, Vector Labs), and the SG peroxidase substrate (Vector Labs). In three rats brain stem sections were first immunoreacted with a goat anti-choline acetyltransferase antibody (1:1000; Millipore Sigma) and a rabbit anti-goat IgG conjugated to horseradish peroxidase (Vector Labs), which was revealed with diaminobenzidine (brown reaction product). Sections were then immunoreacted with a mouse anti-GFP antibody (1:2000; Abcam), a biotinylated anti-mouse IgG, which was revealed with streptavidin horseradish peroxidase conjugate and the Ni-DAB substrate (black reaction product). Finally, the extent of electrolytic lesions was assessed on material stained with Neutral Red. Sections were scanned at a resolution of 0.5 µm/pixel (TissueScope LE; Huron Digital Pathology) and imported in Fiji or Photoshop for color and contrast adjustments.

### Data analysis

Three-dimensional maps of retrogradely labeled KF/PB cells were constructed with the Neurolucida software (Microbrightfield).

Peristimulus time histograms were built using the LabChart 8.0 spike histogram module (AD Instruments), and the Matlab Chronux toolbox (http://www.chronux.org) was used to compute the spectral coherence between respiration and the firing rate of KF/PB cells.

## REFERENCES

Barik, A., Thompson, J.H., Seltzer, M., Ghitani, N., and Chesler, A.T. (2018). A brainstem-spinal circuit controlling nocifensive behavior. Neuron 100, 1491–1503.

Bellavance, M.A., Takatoh, J., Lu, J., Demers, M., Kleinfeld, D., Wang, F., and Deschênes, M. (2017). Parallel inhibitory and excitatory trigemino-facial feedback circuitry for reflexive vibrissa movement. Neuron 95, 673–682.

Bernard, J.F., Huang, G.F., and Besson, J.M. (1992). Nucleus centralis of the amygdala and the globus pallidus ventralis: electrophysiological evidence for an involvement in pain processes. J. Neurophysiol. 68, 551–569.

Bernard, J.F., Villanueva, L., Carroue, J., and Le Bars, D. (1990). Efferent projections from the subnucleus reticularis dorsalis (SRD): a Phaseolus vulgaris leucoagglutinin study in the rat. Neurosci. Lett. 116, 257–262.

Cechetto, D.F., Standaert, D.G, and Saper, C.B. (1985). Spinal and trigeminal dorsal horn projections to the parabrachial nucleus in the rat. J. Comp. Neurol. 240, 153–160.

Chamberlin, N.L. (2004). Functional organization of the parabrachial complex and intertrigeminal region in the control of breathing. Respir. Physiol. Neurobiol. 143, 115–125.

Chamberlin, N.L., and Saper, C.B. (1992). Topographic organization of cardiovascular responses to electrical and glutamate microstimulation of the parabrachial nucleus in the rat. J. Comp. Neurol. 326, 245–262.

Cimadevilla, J.M., Fenton, A.A., and Bures, J. (2001). New spatial cognition tests for mice: passive place avoidance on stable and active place avoidance on rotating arenas. Brain Res. Bull. 54, 559–563.

Curtis, J. C. & Kleinfeld, D. (2009). Phase-to-rate transformations encode touch in cortical neurons of a scanning sensorimotor system. Nat. Neurosci. 12, 492–501.

Dallel, R., Ricard, O., and Raboisson, P. (2004). Organization of parabrachial projections from the spinal trigeminal nucleus oralis: an anterograde tracing study in the rat. J. Comp. Neurol. 470, 181–191.

Dean, P., Redgrave, P., and Westby, G.W.M. (1989). Event or emergency? Two response systems in the mammalian superior colliculus. Trends in Neurosci. 12, 137–147.

Delaney, A.J., Crane, J.W., and Sah, P. (2007). Noradrenaline modulates transmission at a central synapse by a presynaptic mechanism. Neuron 56, 880–892.

Deschênes, M., and Urbain, N. (2016). Vibrissal afferents from trigeminus to cortices. In Scholarpedia of Touch. T.J. Prescott et al. eds. (Atlantis Press) DOI 10.2991/978-94-6239-133-8-49657-672.

Engelmann, M., Thrivikraman, K.V., Su, Y., Nemeroff, C.B., Montkowski, A., Landgraf, R., Holsboer, F., and Plotsky, P.M. (1996). Endocrine and behavioral effects of airpuff-startle in rats. Psychoneuroendocrinol. 21, 391–400.

Feil, K., and Herbert, H. (1995). Topographic organization of spinal and trigeminal somatosensory pathways to the rat parabrachial and Kölliker-Fuse nuclei. J. Comp. Neurol. 353, 506–528.

Frangeul, L., Porrero, C., Garcia-Amado, M., Maimone, B., Maniglier, M., Clascá, F., and Jabaudon, D. (2014). Specific activation of the paralemniscal pathway during nociception. Eur. J. Neurosci. 39, 1455–1464.

Furuta, T., Nakamura, K., and Deschênes, M. (2006). Angular tuning bias of vibrissa-responsive cells in the paralemniscal pathway. J. Neurosci. 26, 10548–10557.

Geerling, J.C., Yokota, S., Rukhadze, I., Roe, D., and Chamberlin, N.L. (2017). Kölliker-Fuse GABAergic and glutamatergic neurons project to distinct targets. J. Comp. Neurol. 525, 1844–1860.

Guo, Z., Li, P., Longhurst, J.C. (2002). Central pathways in the pons and midbrain involved in cardiac sympathoexcitatory reflexes. Neuroscience 113, 435–447.

Igarashi, K.M., Ieki, N., An, M., Yamaguchi, Y., Nagayama, S., Kobayakawa, K., Kobayakawa, R., Tanifuji, M., Sakano, H., Chen, W.R., and Mori.,K. (2012). Parallel mitral and tufted cell pathways route distinct odor information to different targets in the olfactory cortex. J. Neurosci. 32, 7970–7985.

Isett, B. R. & Feldman, D. E. (2020) Cortical coding of whisking phase during surface whisking. Current Biology 30, R934–R936.

Jacquin, M.F., Mooney, R.D., and Rhoades, R.W. (1986). Morphology, response properties, and collateral projections of trigeminothalamic neurons in brainstem subnucleus interpolaris of rat. Exp. Brain Res. 6, 457–468.

Jacquin, M.F., Barcia, M., and Rhoades, R.W. (1989). Structure-function relationships in rat brainstem subnucleus interpolaris: IV. Projection neurons. J, Comp. Neurol. 282, 45–62.

Jansen, N.A., and Giesler, G.J. Jr (2015). Response characteristics of pruriceptive and nociceptive trigeminoparabrachial tract neurons in the rat. J. Neurophysiol. 113, 58–70.

Kleinfeld, D. & Deschênes, M. (2011). Neuronal basis for object location in the vibrissa scanning sensorimotor system. Neuron 72, 455–468.

Knapp, D.J., and Pohorecky, L.A. (1995). An air-puff stimulus method for elicitation of ultrasonic vocalisations in rats. J. Neurosci. Meth. 62, 1–5.

Lara, J.P., Parkes, M.J., Silva-Carvhalo, L., Izzo, P., Dawid-Milner, M.S., and Spyer, K.M. (1994). Cardiovascular and respiratory effects of stimulation of cell bodies of the parabrachial nuclei in the anaesthetized rat. J. Physiol. 477, 321–329.

Lomber, S.G., and Malhotra, S. (2008). Double dissociation of ‘what’ and ‘where’ processing in auditory cortex. Nat. Neurosci. 11, 609–616.

Martins, I., and Tavares, I. (2017). Reticular formation and pain: the past and the future. Front. Neuroanat. 11, 51. doi:10.3389/fnana.00051.

Masri, R., Quiton, R.L., Lucas, J.M., Murray, P.D., Thompson, S.M., and Keller, A. (2009). Zona incerta: A role in central pain. J. Neurophysiol. 102, 181–191.

Merigan, W.H., and Maunsell, J.H. (1993). How parallel are the primate visual pathways? Annu. Rev. Neurosci. 16, 369–402.

Moore, J.D., Mercer L.N., Deschênes, M., and Kleinfeld, D. (2015). Vibrissa self-motion and touch are reliably encoded along the same somatosensory pathway from brainstem through thalamus. PLoS Biol 13(9): e1002253. doi:10.1371/journal.pbio.1002253.

Nassi, J.J., and Callaway, E.M. (2009). Parallel processing strategies of the primate visual system. Nat. Rev. Neurosci. 10, 360–372.

Niu, J., Ding, L., Li, J.J., Kim, H., Liu, J., Li, H., Moberly, A., Badea, T.C., Duncan, I.D., Son, Y.J., Scherer, S.S., and Luo, W. (2013). Modality-based organization of ascending somatosensory axons in the direct dorsal column pathway. J. Neurosci. 33, 17691–17709.

Ogawa H, Nomura T. (1988) Receptive field properties of thalamo-cortical taste relay neurons in the parvicellular part of the posteromedial ventral nucleus in rats. Exp. Brain Res. 73:364–70.

Phelan, K.D., and Falls, W.M. (1991). A comparison of the distribution and morphology of thalamic, cerebellar and spinal projection neurons in rat trigeminal nucleus interpolaris. Neurosci. 40, 497–511.

Pierret, T., Lavallée, P., and Deschênes, M. (2000). Parallel streams for the relay of vibrissal information through thalamic barreloids. J. Neurosci. 20, 7455–7462.

Redgrave, P., Telford, S., Wang, S., McHaffie, J.G., and Stein, B.E. (1996). Functional anatomy of nociceptive neurones in rat superior colliculus. Prog. Brain Res. 107, 403–415.

Rodriguez, E., Sakurai, K., Xu, J., Chen, Y., Toda, K., Zhao, S., Han, B.X., Ryu, D., Yin, H., Liedtke, W, and Wang, F. (2017). A craniofacial-specific monosynaptic circuit enables heightened affective pain. Nat. Neurosci. 20, 1734–1743.

Saper, C.B. (2015). Central autonomic systems. In The rat nervous system, G. Paxinos, ed, (Elsevier Academic Press), pp. 761–815.

Saper, C.B., and Stornetta, R.L. (2014) Central autonomic nervous system. In G. Paxinos (Ed.) The rat nervous system (4th ed.) (pp. 369–388). San Diego: Academic Press.

Severson, K.S., Xu, D., Van de Loo, M., Bai, L., Ginty, D.D., O’Connor D.H. (2017). Active touch and self-motion encoding by Merkel cell-associated afferents. Neuron 94, 666–676.

Slugg, R.M., and Light, A.R. (1994). Spinal cord and trigeminal projections to the pontine parabrachial region in the rat as demonstrated with Phaseolus vulgaris leucoagglutinin. J. Comp. Neurol. 339, 49–61.

Sobolewski, A., Kublik, E., Swiejkowski, D.A., Kaminski, J., and Wrobel, A. (2015). Alertness opens the effective flow of sensory information through rat thalamic posterior nucleus. Eur. J. Neurosci. 41, 1321–1331.

Song, G., Wang, H., Xu, H., and Poon, C.S. (2012). Kölliker-Fuse neurons send collateral projections to multiple hypoxia-activated and nonactivated structures in rat brainstem and spinal cord. Brain Struct. Funct. 217, 835–858.

Takatoh, J., Nelson, A., Bolton, M.M., Ehlers, M.D., Mooney, R., and Wang F. (2013). New modules are added to vibrissal premotor circuitry with the emergence of exploratory whisking. Neuron 77, 346–360.

Takatoh, J., Park, J. H., Lu, J., Li, S., Thompson, P. M., Han, B.-X., Zhao, S., Kleinfeld, D., Friedman, B. & F. Wang F. (2021). Constructing an adult orofacial premotor atlas in Allen Mouse CCF. eLIFE 10, e67291.

Urbain, N., Salin, P.A., Libourel, P.A., Comte, J.C., Gentet, L.J., and Petersen, C.C. (2015). Whisking-related changes in neuronal firing and membrane potential dynamics in the somatosensory thalamus of awake mice. Cell Rep. 13, 647–656.

Veinante, P., Jacquin, M., and Deschênes, M. (2000). Thalamic projections from the whisker sensitive regions of the spinal trigeminal complex in the rat. J. Comp. Neurol. 420, 233–240.

Verhagen JV, Giza BK, Scott TR. (2003) Responses to taste stimulation in the ventroposteromedial nucleus of the thalamus in rats. J Neurophysiol., 89:265–275.

Yu, C., Derdikman D., Haidarliu, S., and Ahissar, E. (2006). Parallel thalamic pathways for whisking and touch signals in the rat. PLoS Biol. 2006; 4: e124 10.1371/journal.pbio.0040124.

Zingg, B., Chou, X.L., Zhang, Z.G., Mesik, L., Liang, F., Tao, H.W., and Zhang, L.I. (2017). AAV-mediated anterograde transsynaptic tagging: mapping corticocollicular input-defined neural pathways for defense behaviors. Neuron 93, 33–47.

